# Alterations in intestinal Proteobacteria and antimicrobial resistance gene burden in individuals administered microbial ecosystem therapeutic (MET-2) for recurrent *Clostridioides difficile* infection

**DOI:** 10.1101/2022.11.04.515081

**Authors:** Ashley M. Rooney, Kyla Cochrane, Stephanie Fedsin, Samantha Yao, Shaista Anwer, Satyender Dehmiwal, Susy Hota, Susan Poutanen, Emma Allen-Vercoe, Bryan Coburn, the MTOP Investigators

## Abstract

Intestinal colonisation with pathogens and antimicrobial resistant organisms (AROs) is associated with increased risk of infection. Fecal microbiota transplant (FMT) has successfully been used to cure recurrent *Clostridioides difficile* infection (rCDI) and to decolonise intestinal AROs. However, FMT has significant practical barriers to implementation. A microbial consortium, microbial ecosystem therapeutic (MET)-2, is an alternative to FMT for the treatment of rCDI. It is unknown whether MET-2 is associated with decreases in pathogens and antimicrobial resistance genes (ARGs). We conducted a post-hoc metagenomic analysis of stool collected from two interventional studies of MET-2 (published) and FMT (unpublished) for rCDI treatment to understand if MET-2 had similar effects to FMT for decreasing pathogens and ARGs as well as increasing anaerobes. Patients were included in the current study if baseline stool had Proteobacteria relative abundance ≥10% by metagenomic sequencing. We assessed pre- and post-treatment Proteobacteria, obligate anaerobe and butyrate-producer relative abundances and total ARGs. MET-2 and FMT were associated with decreases in Proteobacteria relative abundance as well as increases in obligate anaerobe and butyrate-producer relative abundances. The microbiota response remained stable over 4 or 6 months for MET-2 and FMT, respectively. MET-2, but not FMT, was associated with a decrease in the total number of ARGs. MET-2 is a potential therapeutic strategy for ARO/ARG decolonisation and anaerobe repletion.

## Introduction

The human gut microbiome has been implicated as a potential source of infection among hospitalized individuals [1,2]. Gastrointestinal carriage of pathogenic antimicrobial resistant organisms (AROs) with increased abundance of Proteobacteria and decreased abundance of obligate anaerobes and butyrate-producers are associated with elevated risks of infection in these individuals [3–5]. Gut commensal anaerobes provide colonisation resistance against opportunistic pathogens and are important mediators of immune system function and regulation [6]. Butyrate, an anaerobic by-product of dietary fibre fermentation, limits the overgrowth and translocation of opportunistic pathogens by promoting epithelial barrier function, intestinal hypoxia, macrophage antimicrobial function, and immune system homeostasis [6–9].

Antibiotic use, including prophylaxis and selective decontamination of the digestive tract, are effective infection prevention strategies in some populations [5,10]. However, antibiotic use is associated with toxicity, disruption of the gut microbiota, and the growth promotion of AROs [11–13]. Thus, alternatives to antibiotics for infection prevention and control are needed.

Fecal microbiota transplant (FMT) is effective for the treatment of recurrent *Clostridioides difficile* infection (rCDI) and is associated with decreased incidence of bloodstream infection in this population [14,15]. FMT may be a potential strategy for decolonising intestinal AROs where eradication rates have ranged from 37.5 – 87.5% in mostly small observational studies lacking a placebo control [16]. However, FMT is limited by safety and scalability challenges [17,18]. Microbial ecosystem therapeutic (MET)-2, is a bacterial consortium of 39 isolated organisms and has been demonstrated in a recent clinical trial to cure rCDI in 79% of trial participants [19]. Since patients with rCDI are often colonized with AROs and have increased abundances of Proteobacteria including clinically-relevant members of Enterobacteriaceae in the gut microbiome [16,20], we postulated that MET-2 administration may have similar effects to FMT for decreasing Proteobacteria and antimicrobial resistant gene (ARG) abundance in stool.

In this study, we performed a post-hoc metagenomic analysis of stool microbiome composition in recipients of MET-2 or FMT for rCDI from two cohorts to assess their effects on the abundances of Proteobacteria, ARGs, obligate anaerobes, and butyrate-producers in pre- and post-therapy stool samples.

## Methods

### Study design and participants

This is a study of patients 18 years or older with rCDI, defined as one or more recurrences of CDI, who participated in separate prospective cohorts evaluating the effects of either a microbial consortium (MET-2), which has been previously described [19], or FMT (unpublished) on rCDI recurrence. As part of the Microbiota Therapeutics Outcomes Program, the FMT for rCDI study is currently ongoing in Toronto, Ontario, Canada. Donor screening protocol, the preparation of the FMT, as well as participant inclusion and exclusion criteria are outlined in Supplementary Methods 1.

Briefly, both cohorts were on antibiotic therapies prior to the therapeutic intervention. Patients who received FMT underwent bowel preparation prior to FMT. FMT was administered via enema 3 times over the course of 7 days. Patients receiving MET-2 did not receive bowel preparation prior to the intervention. Initially, 10 MET-2 capsules were taken orally for 2 days, and then 3 capsules were taken orally for 8 days. In the current study, we selected individuals from either cohort with ≥10% Proteobacteria relative abundance in the baseline stool sample based on metagenomic analyses and did not receive additional MET-2 or FMT. This study had research ethics approval.

### Sample collection and processing

Stool sample collection from the MET-2 study was described previously [19]. Briefly, stool samples were collected at baseline prior to MET-2, as well as approximately 2 weeks, 1 month, and 4 months post-MET-2. For the FMT study, stool samples were collected at baseline prior to FMT, as well as 1, 3, and 6 months post-FMT. All stool samples were stored at -80°C until further use. Stool samples (0.25 g) were subject to DNA extraction using the DNeasy PowerSoil Pro Kit (Qiagen) and stored at -20°C. DNA concentration was measured using the Qubit Fluorometer (Thermo Fisher) following the manufacturer’s instructions. Prior to library preparation, DNA was diluted to approximately 100 ng in DNase/RNase free water. Sequencing libraries were generated using the DNA Prep kit (Illumina) and the IDT for Illumina UD indexes (Illumina). Libraries were stored at -20°C. Libraries were manually pooled and sequenced at 2 x150 bp using the SP flowcell on the NovaSeq 6000 at the Princess Margret Genomics Centre.

### Outcomes

The primary outcome was change in Proteobacteria relative abundance between baseline and approximately 1-month (30 days ± 10 days) post-intervention (MET-2 and FMT). The secondary outcomes included total ARGs as well as obligate anaerobe and butyrate-producer relative abundance between baseline and 1-month post-intervention (MET-2 and FMT). Lastly, exploratory longitudinal analyses of Proteobacteria, obligate anaerobe, and butyrate-producer relative abundances up to 4 months in the MET-2 interventional group and 6 months in the FMT interventional group were performed.

### Sequence data processing

Sequence quality was assessed with FastQC v.0.11.9 [21]. As the quality was high, no sequence trimming was performed. Nextera adapters were trimmed with Trimmomatic v0.39 [22]. Human and phiX reads were removed with KneadData v.0.7.2 [23]. Taxa were identified from quality-processed reads using Metaphlan3 v.3.0.13 [24]. To ensure an even sampling depth for ARG detection, quality processed-reads were subsampled to 12,328,297 reads, which represents the lowest sequencing depth that retained all baseline and 1-month samples from the MET-2 and FMT interventional groups. Based on the performance characteristics of sequencing metagenomic samples for the detection of ARGs [25], 12,328,297 reads per sample provides a detection frequency ≥90% for all ARGs to relative abundances of ≥3%. The subsampled reads were assembled into contigs using metaSPades v.3.15.3 [26] with the recommended kmer lengths of 21, 33, 55, and 77. To predict the ARGs from metagenome-assembled contigs, RGI *main* v.5.1.0 of the CARD [27] was used on default settings (perfect and strict hits identified only), specifying DIAMOND v.0.8.36 [28] as the local aligner and the *–low_quality* flag. RGI’s *heatmap* v.5.1.0 function was used to categorize ARGs based on drug class-associated resistance. Published 16S rRNA sequences of stool samples from vancomycin-treated patients with CDI pre-treatment and approximately 1-month post-treatment were analyzed to measure Proteobacteria relative abundance, as a reference [29,30], using QIMME2 v.2022.8 [31].

### Proteobacteria quantitative polymerase chain reaction (qPCR) for absolute abundance quantification

Density of γ-Proteobacteria in each fecal sample and DNA extraction negative controls were measured using qPCR, with the forward primer (5’-TCGTCAGCTCGTGTYGTGA-3’), the reverse primer (5’-CGTAAGGGCCATGATG-3’)[32] and probe (HEX-5’-AACGAGCGC-ZEN-AACCCTTWTCCY-3’-FQ-IABk) (Integrated DNA Technologies).

### Microbiota analyses and anaerobe classification

Proteobacteria content in each sample was summarized at the species level and overall relative abundance was quantified at the phylum level. Enterobacteriaceae, γ-Proteobacteria relative abundances were also quantified. Obligate anaerobe and butyrate-producer diversity in each sample was summarized and relative abundance determined at the species level. We used Bergey’s Manual of Systematic Bacteriology volumes 2 – 5 to manually classify species-level taxa as obligate anaerobes and butyrate-producers based on descriptors in the manuals such as “strictly anaerobic”, “anaerobic” or “obligate anaerobe” as well as “produces butyrate”, “forms butyrate”, or “butyric acid is an end product” [33–37]. If the manual did not have descriptive terms for butyrate production, taxa were not considered butyrate-producers. Microbiome measurements were assessed at baseline, 30 days (± 10 days) and up to 4 or 6 months post-intervention for MET-2 and FMT recipients, respectively.

### Statistical analyses

Samples were grouped by intervention received (MET-2 or FMT) and stratified by timepoint. To compare relative abundances between baseline and 1-month post-intervention samples, 0.000001 was added to relative abundances for all taxa (to account for zeros), then log transformed. Pairwise analyses were performed using the non-parametric Wilcoxon matched-pairs signed-rank test on log-transformed relative abundances and total number of ARGs within interventional groups between the baseline and 1-month post-intervention timepoints. The non-parametric Mann-Whitney U-Test was used to compare groups. The non-parametric Spearman’s correlation was performed to test the relationship between total ARGs and Proteobacteria abundance. Statistical analyses were performed in GraphPad Prism v.9.0.3.

## Results

### Participant characteristics

A total 15/19 (79%) MET-2 for rCDI trial participants and 5/8 (62%) FMT for rCDI study participants were included in the current study. Participants in both groups initially received vancomycin, except for participant #1 in the FMT study who received fidaxomicin. The median age of the MET-2 and FMT participants was 65 years and 67 years, respectively, and female patients were more common in both groups (MET-2: 67%, 10/15, FMT: 100% (5/5). A single FMT donor provided stool on multiple dates for preparation of FMT. Supplementary Table 1 provides stool donation dates and to which recipients the donated stool was administered. Participant #10 who received MET-2 failed initial MET-2 administration and was retreated (stool samples before and after re-treatment are not included in this study). Participant #3 of the FMT group failed the initial course of FMT and did not receive another course within the study period. For the baseline and 1-month post-intervention analyses, data from 2 patients from the MET-2 interventional group were excluded due to missing 1-month samples. These 2 patients were included for the longitudinal analyses.

### Proteobacteria abundance pre- and post-intervention

Among included patients, median baseline Proteobacteria relative abundance for participants who received MET-2 (n = 13) was 44% (range, 11% - 97%) and for participants who received FMT (n = 5) was 55% (range, 17% - 93%) (Mann-Whitney p-value: 0.70). At baseline, the most common and abundant Proteobacteria genera in the MET-2 and FMT groups were *Klebsiella* which included *K. pneumoniae, K. oxytoca, K. variicola, K. michiganensis*, and *K. quasipneumoniae*. Other species included *Escherichia coli, Enterobacter cloacae complex, and Citrobacter* spp. (Figure 1A).

**Figure 1.**
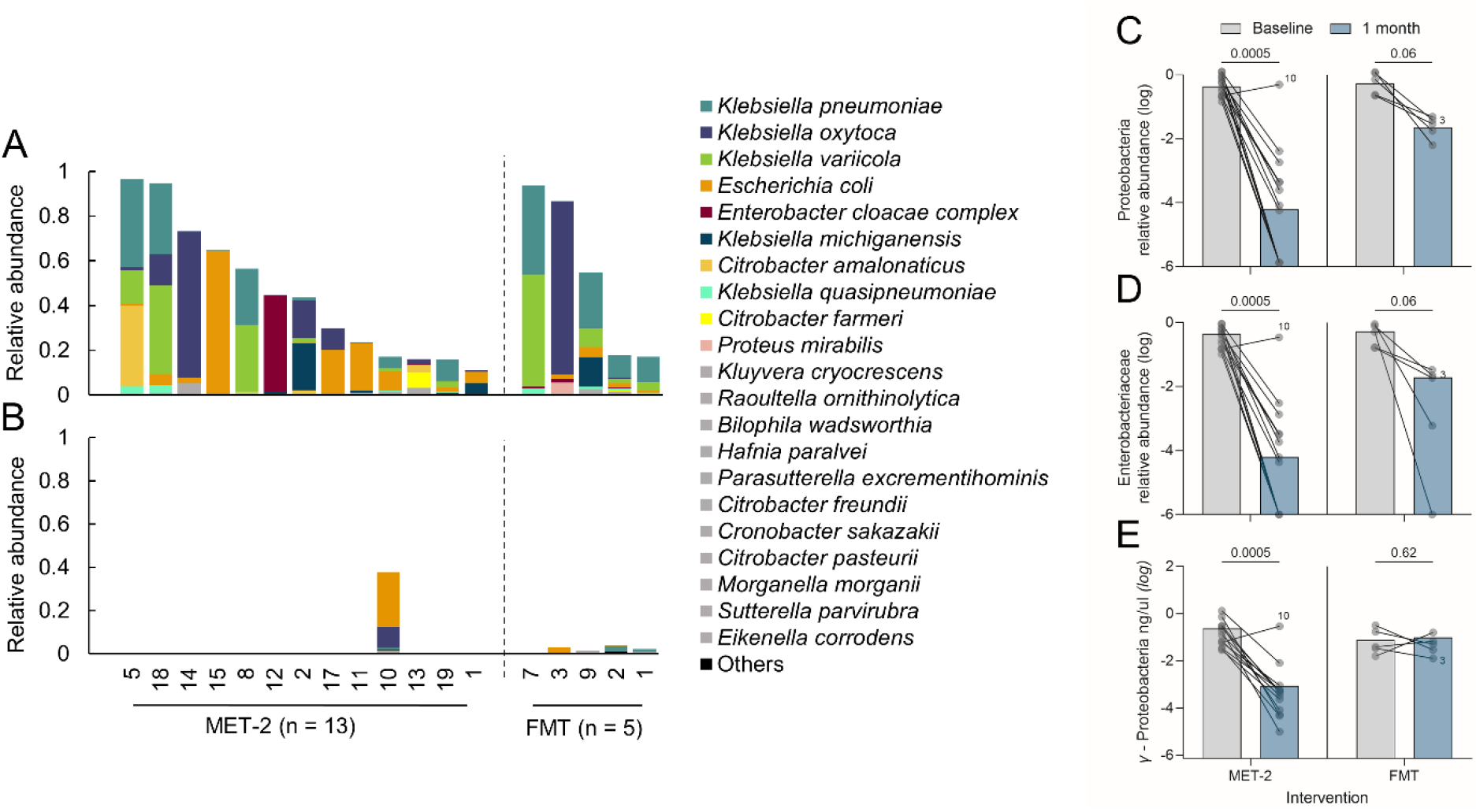
Proteobacteria relative abundances in participants who received MET-2 (n = 13) or FMT (n = 5). Proteobacteria are summarized at the species-level in the baseline (**A**) and 1-month post-intervention stool samples (**B**). **A-B**, Species contributing <1% relative abundance in at least one sample were aggregated as “Others”. Species contributing <5% relative abundance, but ≥1% in at least one sample are in grey. **C**, The log-scale Proteobacteria relative abundance between baseline and 1-month post-intervention. **D**, The log-scale Enterobacteriaceae relative abundance between baseline and 1-month post-intervention. **E**, The log-scale γ-Proteobacteria absolute abundance (ng/μl, limit of detection (log): -6.14 ng/ul) between baseline and 1-month post-intervention. Dots represent individual patients with lines connecting the same patients measured at different time points. Participant 10 and participant 3 are highlighted as individuals who failed initial MET-2 or FMT therapy, respectively. Medians are plotted with p-values displayed above each interventional group. Pairwise analysis performed using Wilcoxon matched-pairs signed rank test.

At 1-month post-intervention, the Proteobacteria relative abundance decreased (Figure 1B), to a median relative abundance of 0.01% (range, 0% - 38%) in the MET-2 group and 2.2% (range, 0.5% - 3.7%) in the FMT group. Between baseline and 1-month post-MET-2/FMT there was a decrease in the relative abundances of Proteobacteria (Figure 1C), γ-Proteobacteria (Supplementary Figure 1), and Enterobacteriaceae (Figure 1D) (MET-2; p-values = 0.0005, FMT; p-values = 0.06) with a median log_2_ fold Proteobacteria decrease of 11.8 and 4.9, respectively. The absolute γ-Proteobacteria abundance decreased between baseline and 1-month post-MET-2 (p-value = 0.006) but did not decrease post-FMT (p-value = 0.62) (Figure 1E). One participant (participant 10), failed initial MET-2 therapy for rCDI [19] and was observed in the current study to have a baseline Proteobacteria relative abundance of 17% that increased to 38% by approximately 1 month post-MET-2 administration.

As a comparator to non-microbial therapies, a total of 6 individuals with CDI treated with vancomycin were identified from published datasets [29,30]. The baseline Proteobacteria abundance was 36% (range, 0.6% - 81%) where the most abundant family was Enterobacteriaceae (Supplementary Figure 2A). At 1 month post-vancomycin treatment, Proteobacteria relative abundance was 38% (range, 14% - 78%)(Supplementary Figure 2B). There was no observed decrease in Proteobacteria or Enterobacteriaceae relative abundance between baseline and 1-month post-vancomycin stool samples (Supplementary Figure 2C & D).

### Antimicrobial resistance genes pre- and post-intervention

Total ARGs were measured between baseline and 1-month post-intervention. The ARG numbers at baseline were similar between interventions (Mann-Whitney p-value: 0.16), where the median number of ARGs for participants who received MET-2 was 78 (range, 46 – 131) and for participants who received FMT was 91 (range, 34 – 104). There was an observed decrease in the total number of ARGs by 1 month after the MET-2 intervention, except for patient 10 where the baseline ARG number was 90 and increased to 99 (Figure 2A). FMT was not associated with a decrease in ARGs within 1-month of administration (Figure 2A). There was a strong positive correlation (Spearman r = 0.70, p <0.0001) between the total number of ARGs detected and Proteobacteria relative abundance in the MET-2 interventional group (Supplementary Figure 3A), while there was a weak positive correlation (Spearman r = 0.36, p = 0.31) between the total number of ARGs detected and Proteobacteria relative abundance in the FMT interventional group (Supplementary Figure 3B).

**Figure 2.**
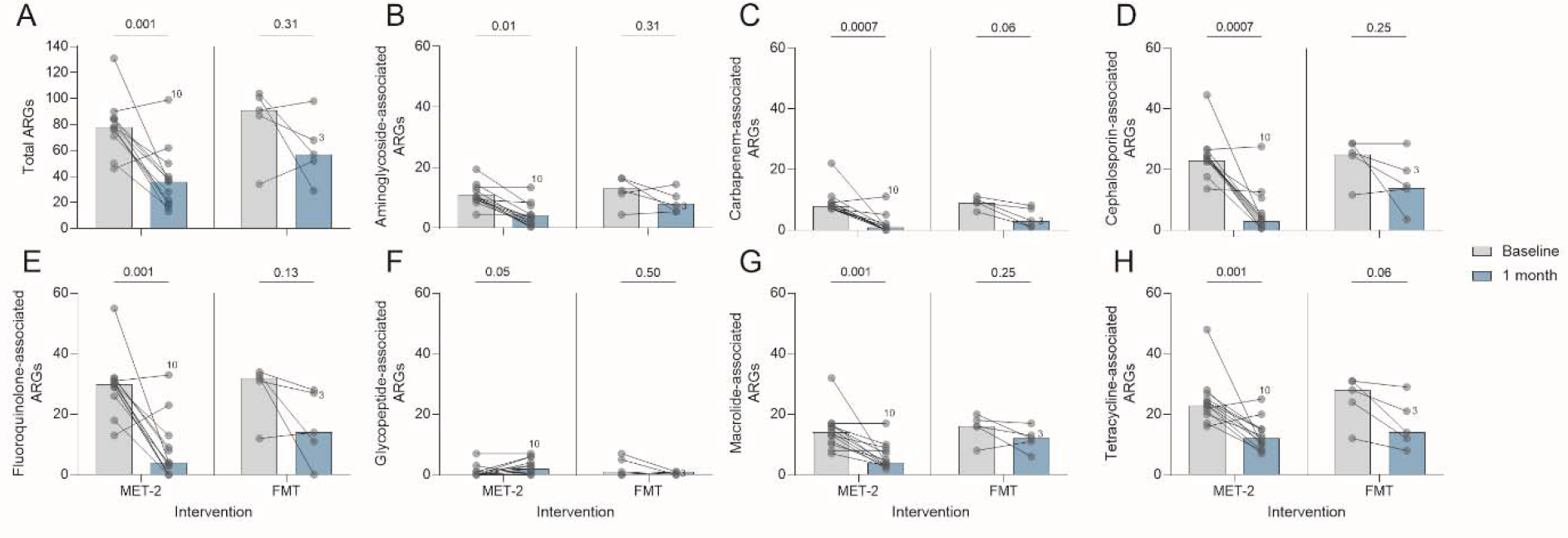
**A**, Antimicrobial resistance genes (ARGs) in participants who received MET-2 (n = 13) or FMT (n = 5) between baseline and 1-month post-intervention. **B-H**, ARGs categorized by drug class-associated resistance in participants who received MET-2 (n = 13) or FMT (n = 5) between baseline and 1-month post-intervention. Drug classes analyzed include aminoglycosides (**B**), carbapenems (**C**), cephalosporins (**D**), fluoroquinolones (**E**), glycopeptides (**F**), macrolides (**G**), and tetracyclines (**H**). Dots represent individual patients with lines connecting the same patients measured at different time points. Participant 10 and participant 3 are highlighted as individuals who failed initial MET-2 or FMT therapy, respectively. Medians are plotted with p-values displayed above each interventional group. Pairwise analysis performed using Wilcoxon matched-pairs signed rank test. and p-values are plotted. Each dot represents an individual with the baseline and 1-month time points included.

The number of ARGs categorized by drug class-associated resistance for the baseline and 1-month post-MET2 or FMT interventions are shown in Figure 2B-H. At baseline in both interventional groups, ARGs conferring resistance to fluoroquinolones (Figure 2E), cephalosporins (Figure 2D), and tetracyclines (Figure 2H) were the most abundant. The number of ARGs associated with resistance to the drug classes analyzed all decreased after MET-2 therapy with the exception of ARGs conferring resistance to glycopeptide antibiotics (Figure 2F). Clinically relevant vancomycin resistance genes (*vanA* or *vanB*) were not detected in 12/13 patients who received MET-2, while *vanA* was detected in patient 10 at baseline and at 1-month post-MET-2 administration (Supplementary Figure 4A). The median *Enterococcus* relative abundance at 1-month post-MET-2 administration was 0% (range, 0% - 21%), where patient 10 had an *Enterococcus* relative abundance of 21% suggesting that the increase in ARGs conferring resistance to glycopeptides in the MET-2 group was not due to vancomycin-resistant *Enterococcus* (Supplementary Figure 3B). FMT was not associated with a decrease in ARGs associated with any drug class assessed (Figure 3B-H).

**Figure 3.**
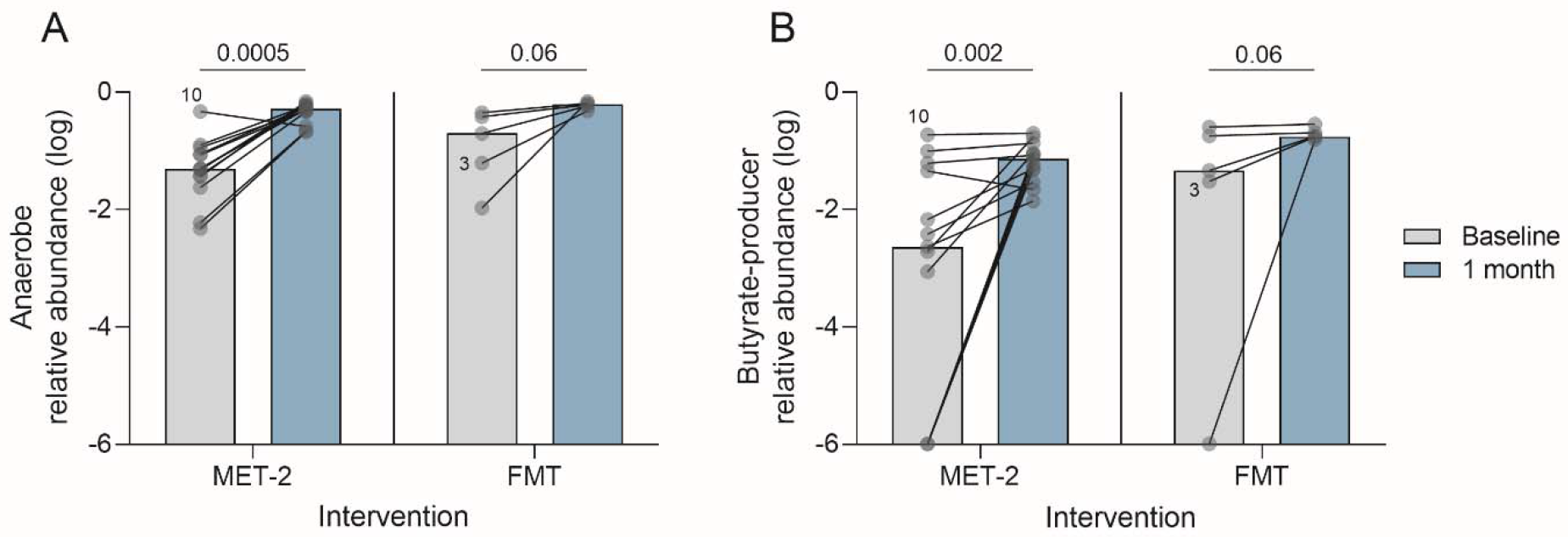
Obligate anaerobes (**A**) and butyrate-producer (**B**) log-scale relative abundances in participants who received MET-2 (n = 13) or FMT (n = 5) between baseline and 1-month post-intervention. Dots represent individual patients with lines connecting the same patients measured at different time points. Participant 10 and participant 3 are highlighted as individuals who failed initial MET-2 therapy. Medians are plotted with p-values displayed above each interventional group. Pairwise analysis performed using Wilcoxon matched-pairs signed rank test.

### Obligate anaerobes and butyrate-producers pre- and post-intervention

Because of their significance in ARO colonisation resistance and intestinal epithelial barrier and systemic immune function, we quantified obligate anaerobes and butyrate-producer relative abundances. At baseline, the median obligate anaerobe relative abundances for the MET-2 and FMT interventional groups were 5%; range, 0.5% - 47% and 20%; range, 1.1% - 45%, respectively (Mann-Whitney p-value: 0.21). The median butyrate-producer relative abundances for MET-2 and FMT interventional groups were 0.002%; range, 0% - 0.2% and 0.05%; range, 0% - 0.3%, respectively (Mann-Whitney p-value: 0.20). There was an observed increase in obligate anaerobe (Figure 3A) (MET-2; p-value = 0.0005, FMT; p-value = 0.06) and butyrate-producer (Figure 3B) (MET-2; p-value = 0.002, FMT; p-value = 0.06) relative abundances between baseline and 1-month post-MET-2 and FMT.

### Microbiota response over time

Proteobacteria, obligate anaerobe, and butyrate-producer abundances were plotted at baseline, 0.5, 1, and 4 months or baseline, 1, 3, and 6 months post intervention for MET-2 (n = 15) and FMT (n = 5), respectively, to assess stability of the treatment-associated changes in microbiome composition. The median Proteobacteria, obligate anaerobe, and butyrate-producer relative abundances remained stable at the final sampling timepoints for both interventions (Figure 4).

**Figure 4.**
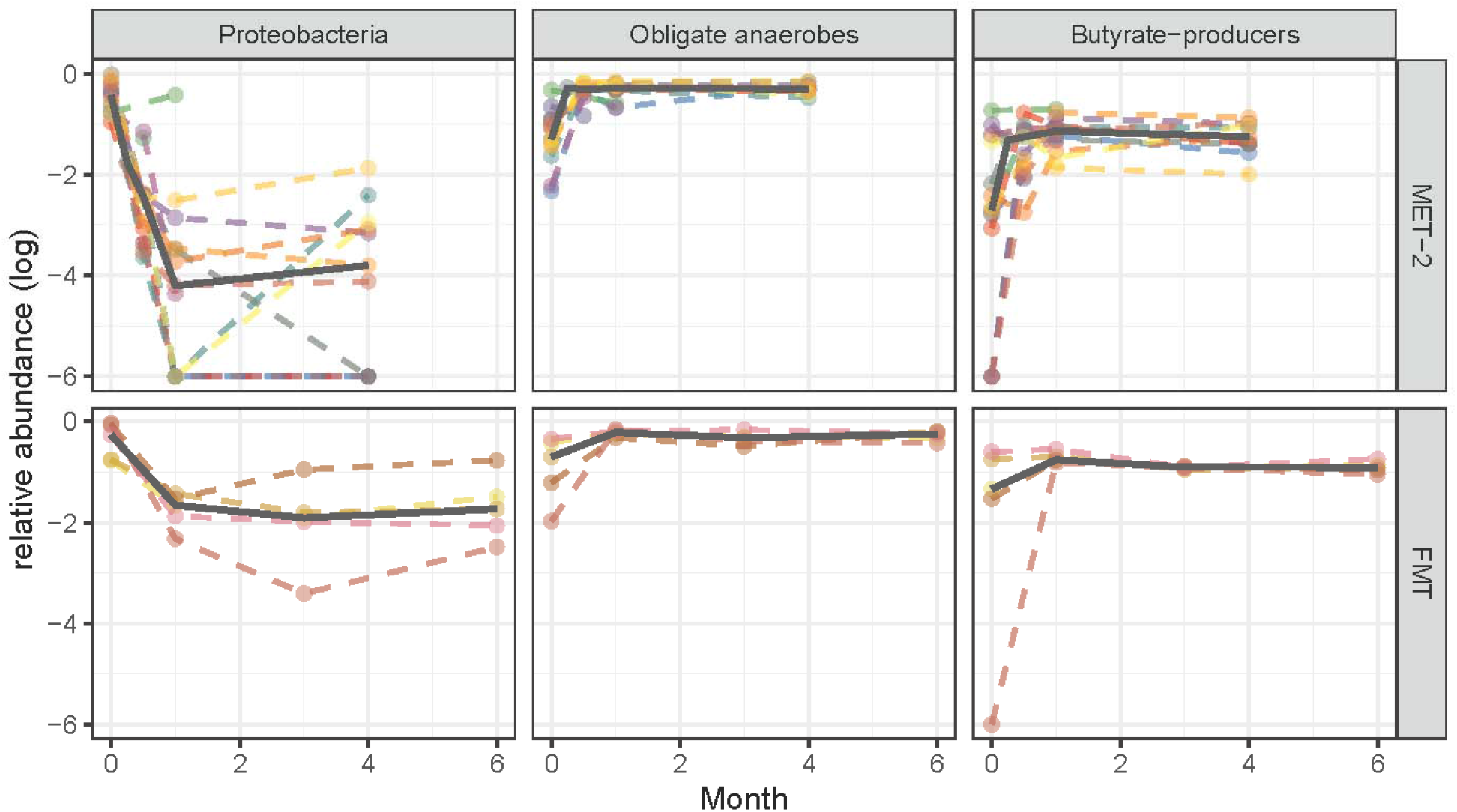
Proteobacteria, obligate anaerobes, and butyrate-producer log-scale relative abundances in participants who received MET-2 (n = 15) or FMT (n = 5) measured over time in months. Dots represent individual patients with dashed lines connecting the same patients measured at different time points. The solid line represents the median across time points.

## Discussion

In the current post-hoc analysis of adult participants with rCDI colonised with Proteobacteria (≥10% relative abundance), we observed that MET-2 had similar effects to FMT for decreasing gut microbiota Proteobacteria relative abundance and increasing obligate anaerobe and butyrate-producer relative abundances by 1-month post-intervention. Absolute abundance of γ-Proteobacteria, measured by qPCR, decreased in the MET-2 interventional group. We did not observe a decrease in the absolute abundance of γ-Proteobacteria in the FMT group suggesting that non-significant trend towards decreasing Proteobacteria relative abundance may not be due to low sample size. Additionally, we observed that the decreased Proteobacteria and increased obligate anaerobes and butyrate-producers observed at 1-month was similar to that at 4 and 6 months for MET-2 and FMT administration, respectively. Lastly, MET-2 and not FMT was associated with decreases in ARG numbers in the gut microbiome by 1-month post-MET-2.

To our knowledge, this is the first study to assess the effects of a therapeutic microbial consortium on Proteobacteria and ARGs in the gut microbiome of patients being treated for rCDI. The results observed in the MET-2 interventional group is similar to other studies using metagenomic sequencing to assess the effects of FMT on the gut microbiota composition of patients with rCDI [38–40]. In a sub-study of an open-label, multicentre, clinical trial of RBX2660, a liquid suspension of donor stool for rCDI treatment, Langdon and colleagues [40] found that RBX2660 was associated with a decrease in Enterobacteriaceae by 2 months post-therapy with a microbiota response that remained relatively stable until the final time point of approximately 6 months.

Although MET-2 was associated with a large decrease in Proteobacteria relative abundance (log_2_ fold decrease of 11.8 in the absence of rCDI recurrence), Proteobacteria relative abundance decreased below 20-30% in both interventional groups by 1-month post-therapy. Given that relative abundances of 20-30% have been previously associated with risks such as bacteremia in allogeneic hematopoietic cell transplant patients [5] and patients in a long-term acute care hospital [41], it is uncertain whether the larger decreases in Proteobacteria abundance observed in the MET-2 recipients is associated with additional benefit over simply decreasing relative abundance below this risk-associated threshold.

We observed that MET-2 was associated with a decrease in ARGs by 1-month which is similar to published reports using FMT for this indication [38–40]. In our analysis, we did not observe ARG decreases in the 5 patients with rCDI who received FMT. We did not sequence the donor material from the FMT group, so could not ascertain if the ARGs are being introduced by the FMT. However, in previous studies, even after extensive screening of donor stool, FMT administration was the source of an antibiotic resistant *E. coli* bacteremia [17], while Leung and colleagues [39] have reported that FMT may be a source of clinically-relevant ARGs.

Our study has multiple limitations, first this was a post-hoc analysis of two separate cohorts that were designed to test the effects of MET-2 or FMT for rCDI treatment. It would be inappropriate to make direct comparisons between the effects of MET-2 and FMT on our measured outcomes, so we aimed to instead understand if MET-2 had similar effects to FMT. Next, there is no placebo-control group to account for spontaneous decolonisation of Proteobacteria and ARGs. However, our analyses of published datasets of patients with CDI pre- and post-vancomycin therapy suggest that it is possible that Proteobacteria can remain colonized or increase in some cases 1-month after vancomycin. Others have also reported persistence of ARGs in the gut microbiome acquired post-antibiotic therapy for up to 2 years [42]. Our results were of the metagenome only and did not include culture-based measurements of ARO colonisation. Although, we observed a multiple-log decrease in Proteobacteria and ARGs, these results may not be associated with complete decolonisation of the gastrointestinal tract or host. Lastly, we did not link microbiome changes to any clinical outcomes. Interestingly, Ianiro and colleagues [15] found that the incidence of bloodstream infection was lower in patients who received FMT for rCDI compared to patients who received standard antibiotics, and our results are consistent with putative mechanisms by which this may occur, through both decreasing pathogen burden and increasing the number of anaerobes associated with colonisation and infection-resistance.

In conclusion, MET-2 has similar effects to FMT for decreasing intestinal Proteobacteria and ARGs, while increasing obligate anaerobes and beneficial butyrate-producers. Our results observed in the MET-2 interventional group require validation in a large placebo-controlled prospective trial that includes outcomes of clinical significance.

## Acknowledgments

We thank and acknowledge the investigators of the Microbiota Therapeutics Outcomes Program (MTOP) Johane Allard, Kenneth Croitoru, Herbert Gaisano, David Guttman, Valerie Taylor, Dana Philpott, and Dan Winer.

## Funding Source

This work was supported by a grant to BC from the Weston Foundation and infrastructure support to BC from the Canadian Foundation for Innovation. MTOP initially received seed funding from a University of Toronto Department of Medicine Integrating Challenge Grant.

## Conflicts of Interest

EAV is a cofounder of Nubiyota and KC is employed by Nubiyota. SH was an investigator in a clinical trial by Finch Therapeutics, for which she received a research grant.

## References

1. Taur Y, Pamer EG. The Intestinal Microbiota and Susceptibility to Infection in Immunocompromised Patients. Curr Opin Infect Dis. 2013;26(4):332.

2. Freedberg DE, Zhou MJ, Cohen ME, Annavajhala MK, Khan S, Moscoso DI, et al. Pathogen colonization of the gastrointestinal microbiome at intensive care unit admission and risk for subsequent death or infection. Intensive Care Med. 2018;44:1203–11.

3. McConville TH, Sullivan SB, Gomez-Simmonds A, Whittier S, Uhlemann AC. Carbapenem-resistant Enterobacteriaceae colonization (CRE) and subsequent risk of infection and 90-day mortality in critically ill patients, an observational study. PLoS One. 2017;12(10):e0186195.

4. Magruder M, Sholi AN, Gong C, Zhang L, Edusei E, Huang J, et al. Gut uropathogen abundance is a risk factor for development of bacteriuria and urinary tract infection. Nat Commun. 2019;10:5521.

5. Stoma I, Littmann ER, Peled JU, Giralt S, van den Brink MRM, Pamer EG, et al. Compositional Flux Within the Intestinal Microbiota and Risk for Bloodstream Infection With Gram-negative Bacteria. Clin Infect Dis. 2021;73(11):e4627–35.

6. Belkaid Y, Hand T. Role of the microbiota in immunity and inflammation. Cell. 2014;157(1):121–41.

7. Kelly CJ, Zheng L, Campbell EL, Saeedi B, Scholz CC, Bayless AJ, et al. Crosstalk between microbiota-derived short-chain fatty acids and intestinal epithelial HIF augments tissue barrier function. Cell Host Microbe. 2015;17:662–71.

8. Byndloss MX, Olsan EE, Rivera-Chávez F, Tiffany CR, Cevallos SA, Lokken KL, et al. Microbiota-activated PPAR-γ-signaling inhibits dysbiotic Enterobacteriaceae expansion. Science (80-). 2017;357(6351):570–5.

9. Schulthess J, Pandey S, Capitani M, Rue-Albrecht KC, Arnold I, Franchini F, et al. The Short Chain Fatty Acid Butyrate Imprints an Antimicrobial Program in Macrophages. Immunity. 2019;50:432–45.

10. Evans L, Rhodes A, Alhazzani W, Antonelli M, Coopersmith CM, French C, et al. Surviving sepsis campaign: international guidelines for management of sepsis and septic shock 2021. Intensive Care Med. 2021;47(11):1181–247.

11. Branch-Elliman W, O’Brien W, Strymish J, Itani K, Wyatt C, Gupta K. Association of Duration and Type of Surgical Prophylaxis With Antimicrobial-Associated Adverse Events. JAMA Surg. 2019;E1–9.

12. Bell BG, Schellevis F, Stobberingh E, Goossens H, Pringle M. A systematic review and meta-analysis of the effects of antibiotic consumption on antibiotic resistance. BMC Infect Dis. 2014;14(13):1–25.

13. Rooney AM, Timberlake K, Brown KA, Bansal S, Tomlinson C, Lee K-S, et al. Each Additional Day of Antibiotics Is Associated With Lower Gut Anaerobes in Neonatal Intensive Care Unit Patients. Clin Infect Dis. 2020;70(12):2553–60.

14. Baunwall SMD, Lee MM, Eriksen MK, Mullish BH, Marchesi JR, Dahlerup JF, et al. Faecal microbiota transplantation for recurrent Clostridioides difficile infection: An updated systematic review and meta-analysis. EClinicalMedicine. 2020;29–30:100642.

15. Ianiro G, Murri R, Sciumè GD, Impagnatiello M, Masucci L, Ford AC, et al. Incidence of Bloodstream Infections, Length of Hospital Stay, and Survival in Patients With Recurrent Clostridioides difficile Infection Treated With Fecal Microbiota Transplantation or Antibiotics A Prospective Cohort Study. Ann Intern Med. 2019;171(10):695–702.

16. Saha S, Tariq R, Tosh PK, Pardi DS, Khanna S. Faecal microbiota transplantation for eradicating carriage of multidrug-resistant organisms: a systematic review Clinical Microbiology and Infection. 2019;25:958–63.

17. DeFilipp Z, Bloom PP, Torres Soto M, Mansour MK, Sater MRA, Huntley MH, et al. Drug-Resistant E. coli Bacteremia Transmitted by Fecal Microbiota Transplant. N Engl J Med. 2019;381(21):2043–50.

18. Hota SS, McNamara I, Jin R, Kissoon M, Singh S, Poutanen SM. Challenges establishing a multi-purpose fecal microbiota transplantation stool donor program in Toronto, Canada. Off J Assoc Med Microbiol Infect Dis Canada. 2019;4(4):218–26.

19. Kao D, Wong K, Franz R, Cochrane K, Sherriff K, Chui L, et al. The effect of a microbial ecosystem therapeutic (MET-2) on recurrent Clostridioides difficile infection: a phase 1, open-label, single-group trial. Lancet Gastroenterol Hepatol. 2021;6(4):282–91.

20. Seekatz AM, Young VB. Clostridium difficile and the microbiota. J Clin Invest. 2014;124(10):4182–9.

21. Andrews S. FastQC: a quality control tool for high throughput sequence data [Internet]. 2010.

22. Bolger AM, Lohse M, Usadel B. Trimmomatic: a flexible trimmer for Illumina sequence data. Bioinformatics. 2014;30(15):2114–20.

23. Huttenhower C. KneadData [Internet]. 2022.

24. Truong DT, Franzosa EA, Tickle TL, Scholz M, Weingart G, Pasolli E, et al. MetaPhlAn2 for enhanced metagenomic taxonomic profiling. Nat Methods. 2015;12(10):902–3.

25. Rooney AM, Raphenya AR, Melano RG, Seah C, Yee NR, Macfadden DR, et al. Performance Characteristics of Next-Generation Sequencing for the Detection of Antimicrobial Resistance Determinants in Escherichia coli Genomes and Metagenomes. mSystems. 2022;7(3):e00022–22.

26. Nurk S, Meleshko D, Korobeynikov A, Pevzner PA. MetaSPAdes: A new versatile metagenomic assembler. Genome Res. 2017;27:824–34.

27. Alcock BP, Raphenya AR, Lau TT, Tsang KK, Bouchard E, Edalatmand A, et al. CARD 2020: antibiotic resistome surveillance with the comprehensive antibiotic resistance database. Nucleic Acids Res. 2019;48:D517–25.

28. Buchfink B, Xie C, Huson DH. Fast and sensitive protein alignment using DIAMOND. Nat Methods. 2015;12(1):59–60.

29. Zuo T, Wong SH, Lam K, Lui R, Cheung K, Tang W, et al. Bacteriophage transfer during faecal microbiota transplantation in Clostridium difficile infection is associated with treatment outcome. Gut. 2018;67:634–43.

30. Abujamel T, Cadnum JL, Jury LA, Sunkesula VCK, Kundrapu S, Jump RL, et al. Defining the Vulnerable Period for Re-Establishment of Clostridium difficile Colonization after Treatment of C. difficile Infection with Oral Vancomycin or Metronidazole. Paredes-Sabja D, editor. PLoS One. 2013;8(10):e76269.

31. Bolyen E, Rideout JR, Dillon MR, Bokulich NA, Abnet CC, Al-Ghalith GA, et al. Reproducible, interactive, scalable and extensible microbiome data science using QIIME 2. Nat Biotechnol. 2019;37(8):848–57.

32. De Gregoris TB, Aldred N, Clare AS, Grant Burgess J. Improvement of phylum-and class-specific primers for real-time PCR quantification of bacterial taxa. J Microbiol Methods. 2011;86:351–6.

33. Brenner DJ, Kreig NR, Staley JT, Garrity GM, editors. Bergey’s Manual of Systematic Bacteriology. Vol. 2, The Proteobacteria Part B, The Gammaproteobacteria [Internet]. 2nd ed. Springer-Verlag; 2005. LXXXII, 2816.

34. Brenner DJ, Kreig NR, Staley JT, Garrity GM, editors. Bergey’s Manual of Systematic Bacteriology. Vol. 2, The Proteobacteria Part C, The alpha-, beta-, delta-, and epsilonproteobacteria. 2nd ed. New York: Springer-Verlag; 2005. xxviii, 1388.

35. Vos P, Garrity G, Jones D, Krieg NR, Ludwig W, Rainey FA, et al., editors. Bergey’s Manual of Systematic Bacteriology. Vol. 3, The Firmicutes. 2nd ed. New York: Springer-Verlag; 2009. xxvi, 1450.

36. Krieg NR, Ludwig W, Whitman W, Hedlund BP, Paster BJ, Staley JT, et al., editors. Bergey’s Manual of Systematic Bacteriology. Vol. 4, The Bacteroidetes, Spirochaetes, Tenericutes (Mollicutes), Acidobacteria, Fibrobacteres, Fusobacteria, Dictyoglomi, Gemmatimonadetes, Lentisphaerae, Verrucomicrobia, Chlamydiae, and Planctomycetes. 2nd ed. New York: Springer-Verlag; 2010. XXVI, 949.

37. Whitman W, Goodfellow M, Kämpfer P, Busse H-J, Trujillo M, Ludwig W, et al., editors. Bergey’s Manual of Systematic Bacteriology. Vol. 5, The Actinobacteria. 2nd ed. New York: Springer-Verlag; 2012. XLVIII, 2083.

38. Millan B, Park H, Hotte N, Mathieu O, Burguiere P, Tompkins TA, et al. Fecal Microbial Transplants Reduce Antibiotic-resistant Genes in Patients With Recurrent Clostridium difficile Infection. Clin Infect Dis. 2016;62(12):1479–86.

39. Leung V, Vincent C, Edens TJ, Miller M, Manges AR. Antimicrobial Resistance Gene Acquisition and Depletion Following Fecal Microbiota Transplantation for Recurrent Clostridium difficile Infection. Clin Infect Dis. 2018;66(3):456–7.

40. Langdon A, Schwartz DJ, Bulow C, Sun X, Hink T, Reske KA, et al. Microbiota restoration reduces antibiotic-resistant bacteria gut colonization in patients with recurrent Clostridioides difficile infection from the open-label PUNCH CD study. Genome Med. 2021;13:28.

41. Shimasaki T, Seekatz A, Bassis C, Rhee Y, Yelin RD, Fogg L, et al. Increased Relative Abundance of Klebsiella pneumoniae Carbapenemase-producing Klebsiella pneumoniae Within the Gut Microbiota Is Associated With Risk of Bloodstream Infection in Long-term Acute Care Hospital Patients. Clin Infect Dis. 2019;68(12):2053–9.

42. Jernberg C, Löfmark S, Edlund C, Jansson JK. Long-term ecological impacts of antibiotic administration on the human intestinal microbiota. ISME J. 2007;1(1):56–66.

