## Supplemental Table 1 for "Alterations in intestinal Proteobacteria and antimicrobial resistance gene burden in individuals administered microbial ecosystem therapeutic (MET-2) for recurrent *Clostridioides difficile* infection"

**Supplementary Table 1.** Donor stool collection and FMT administration dates

| **Date of stool collection** | **Date of FMT administration** | **Recipient #** |
| --- | --- | --- |
| May 11, 2017 | July 17, 2017 | 1 |
| May 11, 2017 | July 19, 2017 | 1 |
| May 11, 2017 | July 21, 2017 | 1 |
| May 11, 2017 | July 25, 2017 | 2 |
| May 16, 2017 | July 27, 2017 | 2 |
| May 17, 2017 | July 31, 2017 | 2 |
| May 17, 2017 | July 17, 2017 | 3 |
| May 17, 2017 | July 19, 2017 | 3 |
| May 22, 2017 | July 21, 2017 | 3 |
| May 30, 2017 | August 21, 2017 | 7 |
| June 1, 2017 | August 23, 2017 | 7 |
| June 1, 2017 | August 25, 2017 | 7 |
| June 6, 2017 | August 21, 2017 | 9 |
| June 11, 2017 | August 23, 2017 | 9 |
| June 12, 2017 | August 25, 2017 | 9 |
