## Supplemental Figures 1 - 4 for "Alterations in intestinal Proteobacteria and antimicrobial resistance gene burden in individuals administered microbial ecosystem therapeutic (MET-2) for recurrent *Clostridioides difficile* infection"

**Supplementary Figures**

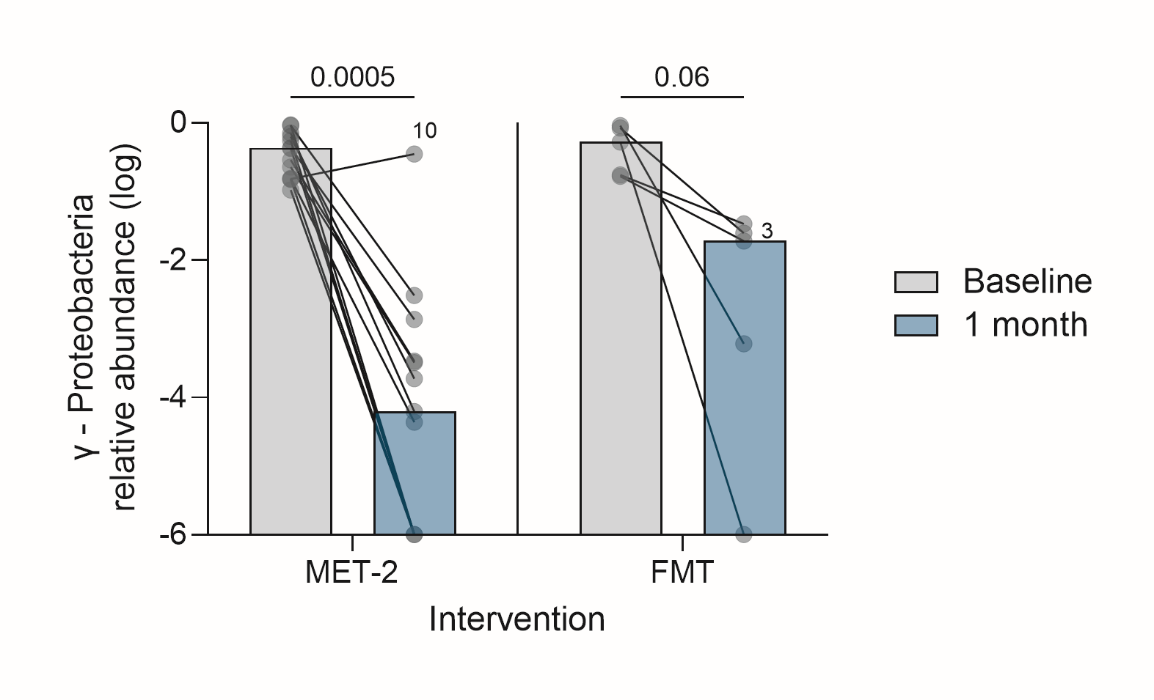

**Supplementary Figure 1.** The log-scale γ-Proteobacteria relative abundance between baseline and 1-month post-intervention. Dots represent individual patients with lines connecting the same patients measured at different time points. Participant 10 and participant 3 are highlighted as individuals who failed initial MET-2 or FMT therapy, respectively. Medians are plotted with p-values displayed above each interventional group. Pairwise analysis performed using Wilcoxon matched-pairs signed rank test.

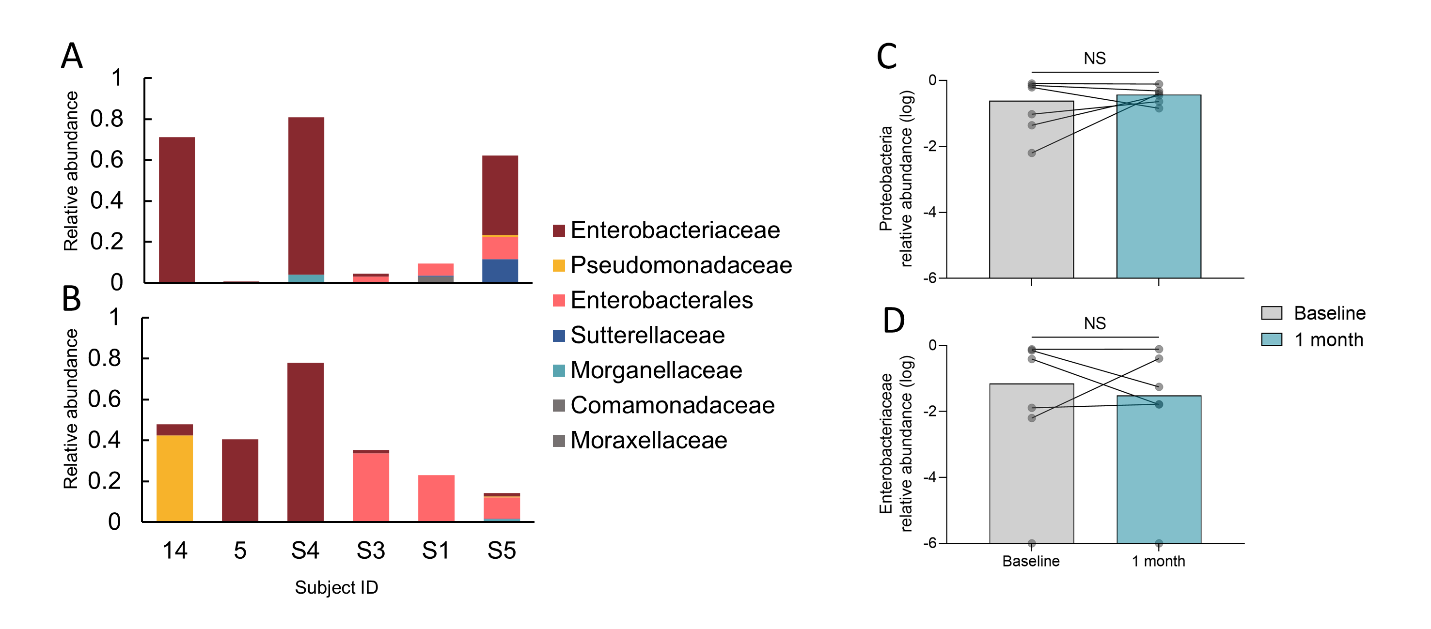

**Supplementary Figure 2.** Histograms of the Proteobacteria relative abundances at baseline (**A**) and 1-month (**B**) post-vancomycin therapy in patients with CDI (n = 6) classified to the family-level. Taxa representing <5% relative abundance are coloured grey. **C,** Log-scale Proteobacteria and (**D**) Enterobacteriaceae relative abundances between baseline and 1-month post-intervention. Dots represent individual patients with lines connecting the same patients measured at different time points. Medians are plotted. Pairwise analysis performed using Wilcoxon matched-pairs signed rank test. P-values were not significant (NS).

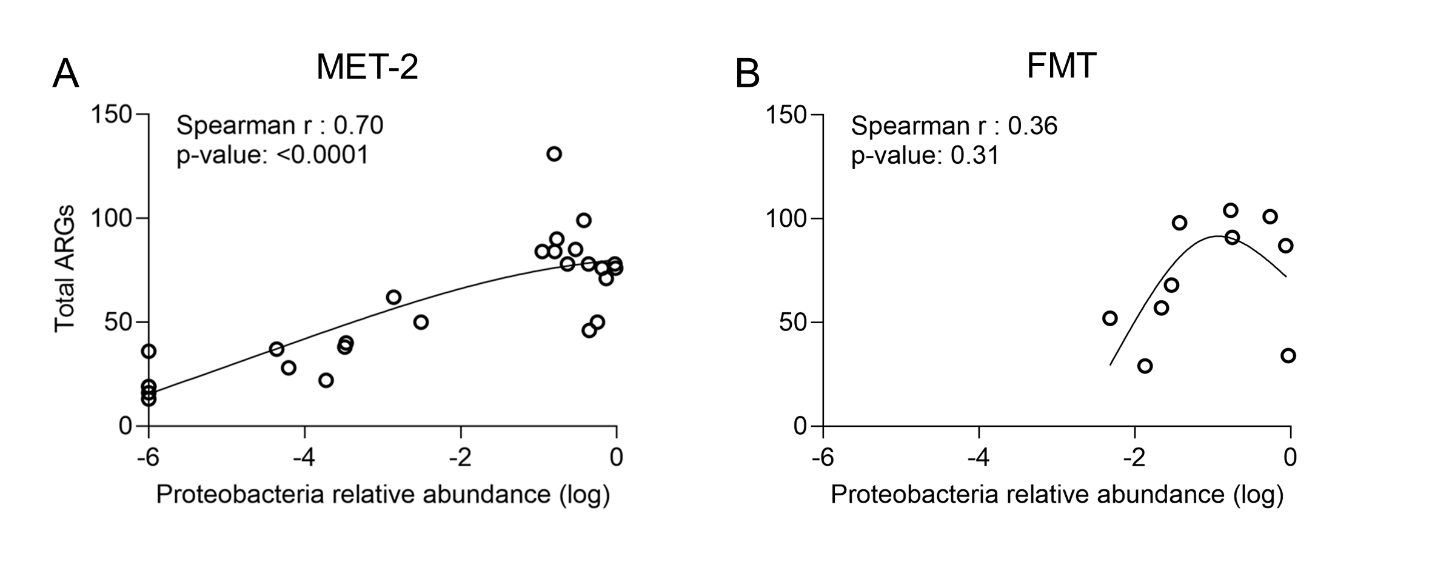

**Supplementary Figure 3. A-B,** The relationship between the number of ARGs and log-scale Proteobacteria relative abundance for individuals who received MET-2 (**A**) and for individuals who received FMT (**B**). **A-B,** Splines are plotted to demonstrate trends. Spearman’s correlation was calculated to measure the relationship between ARGs and log-scale Proteobacteria relative abundance, Spearman’s Rho and p-values are plotted. Each dot represents an individual with the baseline and 1-month time points included.

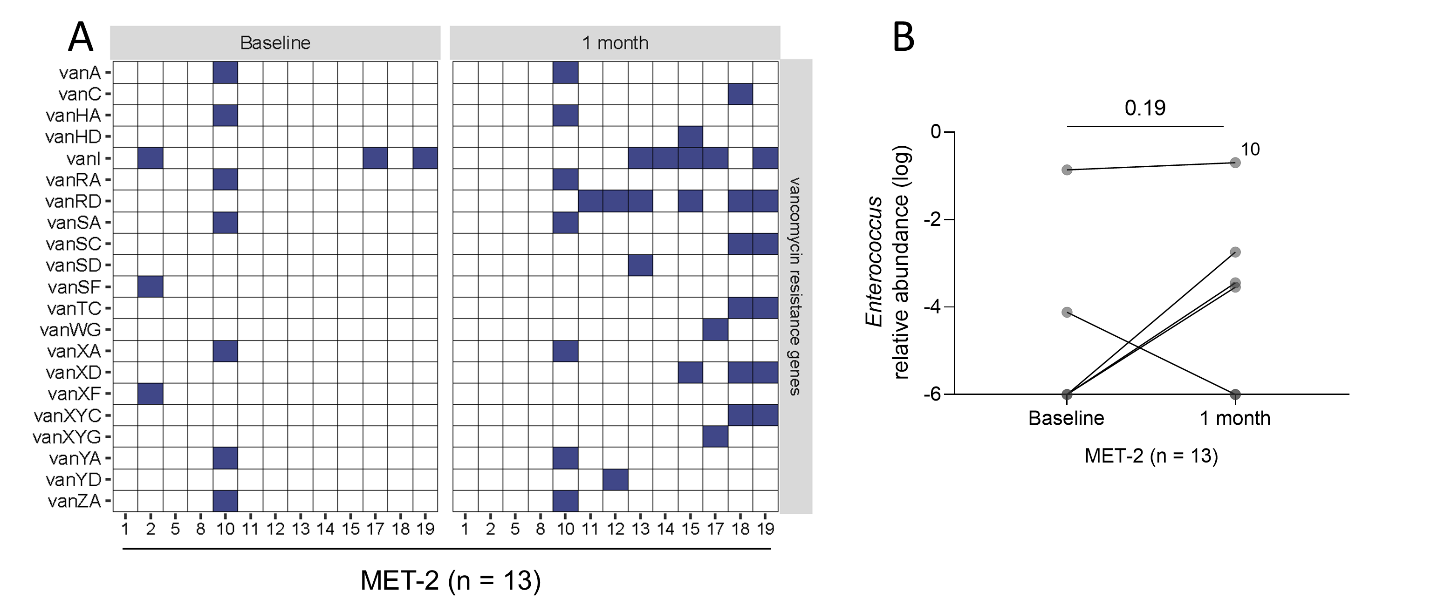

**Supplementary Figure 4. A,** Vancomycin resistance genes detected in stool samples at baseline and 1 month post-MET-2 administration. Blue squares represent presence of a vancomycin resistance gene, while white squares represent absence. **B**, Log-scale *Enterococcus* relative abundance in participants who received MET-2 (n = 13) between baseline and 1-month post-intervention. Dots represent individual patients with lines connecting the same patients measured at different time points. Participant 10 is highlighted as an individual who failed initial MET-2 therapy. Medians are plotted with the p-value displayed above each time point. Pairwise analysis performed using Wilcoxon matched-pairs signed rank test.
