## Supplemental Methods for "Alterations in intestinal Proteobacteria and antimicrobial resistance gene burden in individuals administered microbial ecosystem therapeutic (MET-2) for recurrent *Clostridioides difficile* infection"

**Supplementary Methods**

The following methods are referenced from the protocol: Long-term Measurement of Multidisciplinary Outcomes in Patients Receiving Fecal Microbiota Transplantation (FMT) in the University of Toronto Microbiota Therapeutics Outcomes Program (MTOP) version February 2017.

**FMT Donors**

All FMT donors are considered healthy adults between 18 and 45 years of age. All donors were subject to screening using a self-administered questionnaire for risk-associated behaviors, medical assessment and blood, stool and urine testing for transmissible infectious diseases and mental health questionnaires to exclude mental disorders. This screening is in compliance with the 2015 Health Canada guidance for FMT for recurrent *Clostridioides difficile* infection (rCDI) (1). Full, written informed consent was obtained prior to collecting donor screening or medical assessment data or accepting any donations.

Donors were excluded for the following criteria:

- History or symptoms suggestive of underlying gastrointestinal disease, such as:
  - Inflammatory bowel disease
  - Gastrointestinal motility disorder
  - Diverticular disease
  - Other chronic symptoms of diarrhea NYD
  - Episode of acute diarrhea in the last 3 months
  - Irritable bowel syndrome
  - Chronic constipation
- Infection or colonization with transmissible agents, including:
  - HIV-1/2
  - Hepatitis A, B and C viruses
  - HTLV-1/2
  - Syphilis
  - VRE, MRSA, Carbapenemase-producing Organisms (CPO)
  - *Clostridium difficile*
  - *Salmonella*, *Shigella*,Enterotoxigenic *E. coli*, *E.coli* 0157-H7 and other Shiga-toxin producing *E. coli*, *Yersinia,* *Campylobacter, Aeromonas, Plesiomonas,* and *Vibrio*
  - Enteric viruses (norovirus, rotavirus and adenovirus)
  - Ova and parasites
  - *Helicobacter pylori*
  - Gonorrhea
  - Chlamydia
- History of active malignancy, or any cancer within the last 5 years (excluding basal cell carcinoma of skin).
- Risk factors for prion-related disease, including: family history of Creutzfeld-Jacob Disease, corneal or dural transplant, or receipt of human-derived pituitary growth factor.
- History of high risk for recent acquisition of HIV, Hepatitis B or Hepatitis C, as determined by self-screening using a questionnaire of risk-associated behavior
- History or signs of chronic sexually transmitted infections, such as genital ulcerative diseases, anogenital herpes, anogenital warts, chancroid or syphilitic lesions.
- Immunosuppression, as determined by history or suggestive physical signs. Things on history that indicate immunosuppression include:

o   use of any immunosuppressive medication within last 6 months, including any dose of prednisone

o   chronic underlying condition known to produce immunosuppression (eg. systemic lupus erythematosus, end-stage renal disease)

- Physical signs suggestive of immunosuppression include:

o   oral thrush

o   lesions of Kaposi sarcoma

o   disseminated lymphadenopathy

- History or physical findings of chronic liver disease or cholestasis
- Evidence of active encephalitis or meningitis
- Evidence of active systemic viral, bacterial or fungal infection, including malaria and tuberculosis
- Receipt of live vaccine in preceding 30 days
- Receipt of blood transfusion from a country other than Canada in preceding 12 months
- History of dementia or degenerative neurological disorders of unknown etiology
- Recent bite from an animal that may have rabies within the past 6 months
- Antibiotic use in the 6 months preceding donation
- Probiotic agent use for medicinal purposes in the 3 months prior to donation
- Use of cholestyramine within 3 months of donation
- Known and current history of blood in stools
- Travel outside Canada / United State in the past 6 months
- History, family history in first degree (blood) relatives, or current screening symptoms (as determined by positive MINI questionnaire, CD-RISC resilience, Ham-A, MADRS) of psychiatric illness (including depression, anxiety disorder, post-partum depression, bipolar disorder, schizophrenia)
- Family history in first degree (blood) relatives of colon cancer
- History or family history of autoimmune disease in first degree (blood) relatives
- Anaphylactic food and environmental allergies
- Actual BMI lower than 18.5 or higher than 23 kg/m^2^
- Diabetes or “pre-diabetes”, defined as HbA1c > 6% according to FMT screening protocol, or HOMA-IR more than 2.73
- Overweight (BMI >/= 25) at any time during lifetime
- Waist circumference > 80 cm for women, > 102 cm for men
- Smoker (cigarette, marijuana, other)
- Use of recreational drugs
- Greater than 2 alcoholic beverages per day on a regular basis

**FMT Donor Re-screening Procedures**

As donors provided repeat donations, re-screening was performed as follows. Re-screening for infectious diseases (stool, urine, and blood) occurred every 1-3 months since initial screening. FMT donations up to 2 weeks preceding re-screening were quarantined until such time that re-screening for infectious diseases was completed and was negative. A self-screening questionnaire was administered with each donation to identify development of new health conditions, medications and risk-associated behaviors.

**FMT preparation**

FMT preparation occurred at the clinical microbiology laboratory at University Health Network – Sinai Health System. FMT was prepared based on our previous randomized controlled trial (NCT01226992), with modifications. Briefly, a single FMT dose (50 g of screened donor stool) is thawed and diluted to 45 mL of sterile 0.9 N NaCl + 5 ml glycerol. The diluted stool was then filtered through a sterile 330 micron micro-filter-separated- double-compartment polyethylene bag and homogenized with the Stomacher® Paddle Blender. The filtrate was transferred to a 50 mL screw-cap tube and stored at -80°C. An aliquot was taken for storage at -80°C for further testing if needed.

One Falcon tube of frozen FMT was thawed for approximately 2.5 hours at room temperature and diluted to a final volume of 300 ml using 0.9N NaCl. Diluted FMT was then transferred to an enema bag for a single “FMT dose” by enema.

**FMT recipients**

In the current study, FMT recipients were excluded for the following:

- Patients with conditions in whom enemas or colonoscopy are relatively contraindicated
- Unable to tolerate FMT procedure for any other reason
- Receiving an investigational medication
- Pregnant or breastfeeding
- Serious bleeding disorder, anticoagulant use that cannot be stopped temporarily for FMT procedure (or serious platelet disorder (platelet counts below 50)
- Any condition that would pose a health risk to the subject.

All patients receiving FMT and who donated stool samples for microbiome analyses for the current study provided informed consent.

**References**

1. Guidance document: Fecal microbiota therapy used in the treatment of *Clostridium difficile* infection not responsive to conventional therapies. <http://www.hc-sc.gc.ca/dhp-mps/consultation/biolog/fecal_microbiota-bacterio_fecale-eng.php> (accessed April 16, 2016).
